# Progressive shifts in the gut microbiome reflect prediabetes and diabetes development in a treatment-naive Mexican cohort

**DOI:** 10.1101/710152

**Authors:** Christian Diener, Lourdes Reyes, Lilia Jimenez, Mariana Matus, Claudia Gomez, Nathaniel D. Chu, Vivian Zhong, Elizabeth Tejero, Eric Alm, Osbaldo Resendis-Antonio, Rodolfo Guardado-Mendoza

## Abstract

Type 2 diabetes (T2D) is a global epidemic that affects more than 8% of the world’s population and is a leading cause of death in Mexico. Diet and lifestyle are known to contribute to the onset of T2D. However, the role of the gut microbiome in T2D progression remains uncertain. Associations between microbiome composition and diabetes are confounded by medication use, diet, and obesity. Here we present data on a treatment-naive cohort of 405 Mexican individuals across varying stages of T2D severity. Associations between gut bacteria and more than 200 clinical variables revealed a defined set of bacterial genera that were consistent biomarkers of T2D prevalence and risk. Specifically, gradual increases in blood glucose levels, beta cell dysfunction, and the accumulation of measured T2D risk factors was correlated with the relative abundances of four bacterial genera. In a cohort of 25 individuals, T2D treatment - predominantly metformin - reliably returned the microbiome to the normoglycemic community state. Deep clinical characterization allowed us to broadly control for confounding variables, indicating that these microbiome patterns were independent of common T2D comorbidities, like obesity or cardiovascular disease. Thus, our work provides the first solid evidence for a direct link between the gut microbiome and T2D in a critically high-risk population. Whether or not these T2D-associated changes in the gut contribute to the etiology of T2D or its comorbidities remains to be seen.

## Introduction

Type 2 diabetes (T2D) is an acquired multifactorial disease that affects more than 8% of the worldwide population and leads to insulin resistance and insufficient insulin production by pancreatic islet cells ^1–3^. Disease onset is driven or modulated by a variety of factors such as lifestyle, diet, and genetics ^4–7^. T2D incidence is progressively increasing in the Mexican population and has become a major burden for the national health system and one of the leading causes of death in Mexico ^8–10^. The particular vulnerability of the Mexican population to this disease is driven by common factors such as a sedentary lifestyle and diet but is also influenced by risk factors that are specific to the native population ^11^. For instance, it has been shown that about half of all native Mexicans carry a SLC16A11 variant that increases T2D risk by 20% for each haplotype ^12–14^. Consequently, there is an urgent need for diagnosis and treatment strategies in order to limit the progression of T2D in the Mexican high risk population.

Recently, the gut microbiome has been proposed as an important modulator in the progression of T2D. Several studies have shown a wide array of associations between the gut microbiome and diabetes in European, American and Chinese cohorts ^15,16^. Most of those papers have suggested that the diabetic microbiome is less efficient in producing short chain fatty acids (SCFA) due to a loss of butyrate-producing genera ^17–19^. However, especially when looking across different populations, the bacterial genera associated with diabetes vary ^17^, which is consistent with findings that the gut microbiome composition varies greatly across populations ^20^. For example, an increase of Proteobacteria in T2D was reported for Chinese cohorts but was absent in a European cohort ^15,16^. Finding robust associations between the microbiome and T2D is further confounded by treatment effects and comorbidities. Metformin, one of the most common medications T2D, has been shown to modify the gut microbiome which may contribute to its mechanism of action ^21,22^. Indeed, studies comparing diabetic treatment naive individuals with diabetic metformin-treated individuals showed that most of the associations initially attributed to disease progression were a consequence of the treatment alone and absent in individuals without a metformin treatment history ^23^. Apart from medication, changes in lifestyle or diet may also drive changes in the gut microbiome in a disease-independent manner ^24,25^. Thus, two major treatment regimes for T2D, metformin treatment and lifestyle intervention, will likely both trigger their own changes in the gut microbiome and need to be accounted for. Even when isolating the disease from treatment effects, associations may be confounded by comorbidities. The development of T2D is often linked with obesity, a major risk factor in the development of the disease ^20,26^. Additionally, T2D increases risk for cardiovascular disease, which itself has been linked changes in the gut microbiome ^27,28^. Controlling for all of these factors (disease treatment, lifestyle and diet, and comorbidities) might clarify the true associations between the gut microbiome and T2D disease progression. This requires deep phenotyping of the study participants where one measures not only clinically variables related to the disease of interest but also from other groups such as obesity, cardiovascular health, lifestyle and diet. Even though this strategy has been shown to successful in healthy individuals^29^, very few studies have done so in the context of T2D.

To address these concerns and explore the relationship between the microbiome and T2D in an understudied population, we present a controlled study in a Mexican cohort from a distinct geographical region which was specifically designed to avoid those shortcomings. Except for a small control group, all participants in the study were treatment naive and had never received a prior prediabetes or diabetes diagnosis. We also combined a large array of clinical variables related to diabetes with additional phenotype measurements characterizing the lifestyle, diet, obesity prevalence, and cardiovascular health of each individual. This strategy provided a set of more than 200 clinical variables for each individual, allowing us to control for lifestyle and comorbidities and tease out associations specific to different stages of T2D progression. As a result, we identified a set of four bacterial genera that associated consistently with T2D development. Our work establishes a set of gut microbiome markers for type 2 disease progression in a Mexican population independent of treatment effects or secondary phenotypes.

## Results

### The microbiome of treatment-naive individuals associates with a wide range of clinical variables

We recruited a cohort of treatment-naive subjects from the Guanajuato region of Mexico as part of the CARE-In-DEEP Study (Cardiometabolic Risk Evaluation and Interdisciplinary Diabetes Education and Early Prevention) of the University of Guanajuato. This cohort consisted of 405 individuals with no previous diabetes diagnosis and a control group of 25 subjects with previously diagnosed T2D or a history of metformin treatment (see Fig. 1A). Each of the participants in the study underwent extensive clinical characterization consisting of direct measurements as well as a set of validated questionnaires, forming a data set of 226 clinical variables spanning the areas of diabetes, obesity, general health, lifestyle and diet (Fig. 1B). Based on an oral-glucose tolerance test, subjects were stratified into five metabolic groups ranging from normoglycemia and normal glucose tolerance (NG), to different types of prediabetes (impaired fasting glucose, IFG, impaired glucose tolerance IGT, and IFG+IGT), and T2D (see Methods and Fig. 1C). As shown in Table 1, clinical phenotype varied widely between metabolic groups, with a progressive increase in weight, body fat, glycated hemoglobin (HbA1c), glucose levels, and deteriorating insulin sensitivity and pancreatic beta cell function from the NG group to the T2D group (see Fig. S1 A-C).

**Figure 1.**
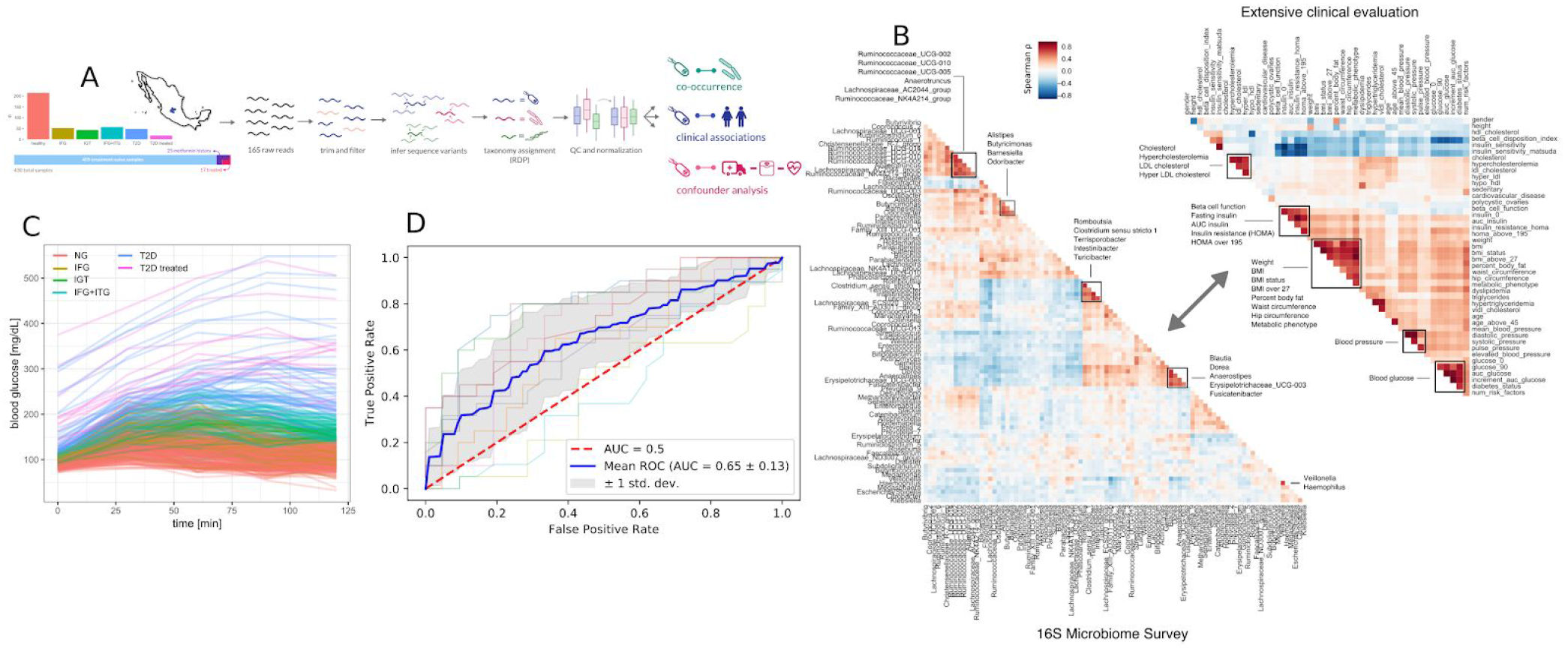
Study design. (A) 405 Individuals were recruited from Guanajuato state and classified in to normoglycemic (NG), impaired fasting glucose (IFG), impaired glucose tolerance (IGT), impaired fasting glucose and impaired glucose tolerance (IFG+IGT), and type 2 diabetes (T2D). 25 individuals under treatment for a previous T2D diagnosis or with previous metformin history were added as controls (T2D treated). (B) Correlations between bacterial genera in the study (intra-microbiome) are shown in the left correlation matrix whereas correlations between clinical variables are shown in the right correlation matrix. (C) Blood glucose curves for all individuals in the study colored by classification. (D) Receiver-Operator curves for predictions from a Random Forest model. Individual cross-validation curves are shown along with the mean trend and standard deviations.

**Table 1.** Cohort characteristics. P value column denotes p values of ANOVA with Bonferroni correction. Superscript letters denote the following: (a) p<0.01 vs NG (b) p<0.01 vs IFG (c) p<0.01 vs IGT (d) p<0.01 vs IFG+IGT (e) p<0.01 vs treatment naive T2D

| Variable | NG | IFG | IGT | IFG+IGT | new T2D | treated T2D | p value |
| --- | --- | --- | --- | --- | --- | --- | --- |
| (n=430) | (n=214) | (n=52) | (n=42) | (n=57) | (n=48) | (n=17) |  |
| **Age (y)** | 38 ± 14 | 45 ± 14<sup>a</sup> | 45 ± 12 | 49 ± 12<sup>a</sup> | 51 ± 9<sup>a</sup> | 55 ± 11<sup>a</sup> | <0.001 |
| **PP(mmHg)** | 38 ± 9 | 41 ± 10 | 42 ± 8 | 45 ± 11<sup>a</sup> | 45 ± 12<sup>a</sup> | 44 ± 12 | <0.001 |
| **Weight (kg)** | 70 ± 15 | 77 ± 17 | 76 ± 15 | 85 ± 17<sup>a</sup> | 85 ± 20<sup>a</sup> | 74 ± 23 | <0.001 |
| **BMI** | 26.5±5.4 | 28.2±5.1 | 29.3±5.3<sup>a</sup> | 32.4±6.2<sup>ab</sup> | 32.5±7.0<sup>ab</sup> | 31.0±9.0<sup>a</sup> | <0.001 |
| **Body fat%** | 33 ± 8 | 32 ± 7 | 35 ± 8 | 39 ± 8<sup>ab</sup> | 39 ± 10<sup>ab</sup> | 37 ± 9 | <0.001 |
| **Visceral fat %** | 6.6 ± 4.1 | 9.3 ± 3.9<sup>a</sup> | 9.3 ± 3.9<sup>a</sup> | 11.8 ± 4.3<sup>a</sup> | 12.2 ± 5.6<sup>a</sup> | 10.3 ± 4.2 | <0.001 |
| **WC (cm)** | 84.7 ±13.4 | 90.3±11.2 | 92.0±10.3<sup>a</sup> | 98.4±13.3<sup>a</sup> | 97.7±18.5<sup>a</sup> | 95.4±10.5 | <0.001 |
| **HbA1c %** | 5.3±0.3 | 5.4±0.3 | 5.4±0.4 | 5.6±0.4 | 6.6±1.6<sup>abcd</sup> | 7.5±1.2<sup>abcd</sup> | <0.001 |
| **Total cholesterol (mg/dl)** | 181 ± 37 | 189 ± 36 | 190 ± 32 | 196 ± 42 | 188 ± 30 | 218 ± 30<sup>a</sup> | <0.001 |
| **TG (mg/dl)** | 131 ± 62 | 173 ± 93<sup>a</sup> | 171 ± 78<sup>a</sup> | 181 ± 80<sup>a</sup> | 200 ± 85<sup>a</sup> | 202 ± 66<sup>a</sup> | <0.001 |

To identify links between the microbiome and the progression of T2D, we sequenced the 16S rDNA gene from stool samples from this cohort. Sequencing data was analyzed using DADA2 which identified 17,059 exact amplicon sequence variants across all samples (see Methods). These sequence variants mapped to 378 bacteria genera, however only 629 sequence variants and 125 genera were appreciably frequent across samples (>10% of individuals). Previous studies have found metformin treatment to lower Intestinibacter abundances and to increase Escherichia abundances ^23^. We found similar trends in our data, albeit not significant (Mann-Whitney p=0.05 and 0.07 for Instestinibacter and Escherichia, see Fig. S1D). In general, T2D could only be weakly predicted from microbiome composition (Random Forest area under ROC = 0.69, see Fig. 1D).

We identified potential links between the microbiome by exhaustive testing of all combinations between bacterial genera and clinical variables, including alpha diversity (Shannon index). Associations between the microbiome and clinical variables were identified by a robust testing strategy based on DESeq2 (see Materials and Methods). Of the 30,780 tests, 208 were deemed significant under an FDR cutoff of 0.05 (Fig. 2A-B). Clinical measurements related to obesity had the most significant associations with microbiome features, while diet-related variables were the least likely to yield a significant association (Fig. 2A). The relative paucity of associations between the microbiome and diet may be a consequence of the heterogeneity of dietary questionnaire data. The genera associating with the most clinical variables was the facultative anaerobe Escherichia and the obligate anaerobe Veillonella, which had 36 and 23 significant associations respectively (Fig. 2B). Escherichia associated mostly with variables related to diabetes and obesity whereas Veillonella associated with variables from many all categories. Ruminococcaceae genera were the most positively correlated with alpha diversity (Shannon index) whereas Fusobacterium, Flavonifractor and Parasutterella were the most negatively associated with alpha diversity (Shannon index, Fig. 2C).

**Figure 2.**
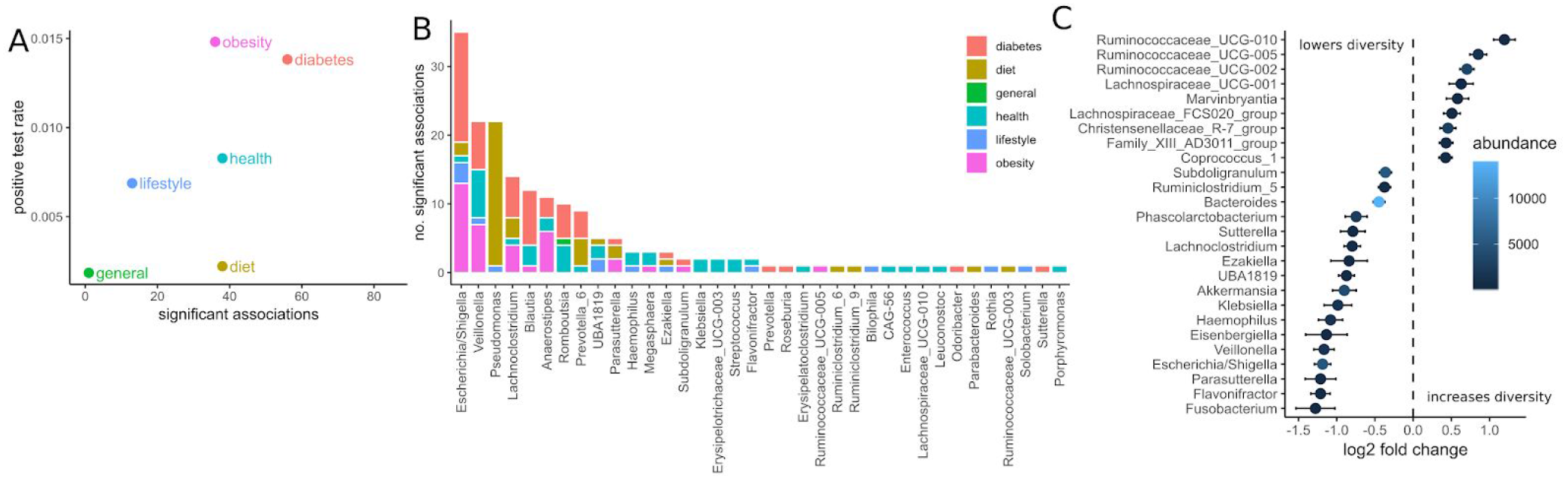
Associations between the microbiome and phenotype. (A) Number of significant associations between microbiome and clinical variables grouped by category (FDR corrected p<0.05). Positive test rate denotes the significant tests / total tests for the category. (B) Significant tests per genus (FDR corrected p<0.05). Color denotes the category of clinical variable the genus associates with. (C) Significant associations (FDR < 0.05) between bacterial genera and alpha diversity (Shannon). Points denote the log fold change (DESeq2 coefficient) of a genus when the diversity increases by one standard deviation. Error bars denote the standard error of the coefficient. Fill color denotes mean no. of normalized reads across all samples.

The gut microbiome of the treatment-naive cohort associated widely with T2D-related clinical variables. A set of 14 bacterial genera associated at least weakly with 25 of the 31 diabetes-related measures (FDR-corrected p-value < 0.05). However, we observed large differences in how those associations distributed across genera (Fig. 3A). Whereas some genera associated with a wide array of T2D measures (for instance Escherichia/Shigella) other associated only with a single measure (e. g. Ezakiella with T2D family history) or exclusively with glucose-related measures, but not insulin-related measures (e. g. Romboutsia, Fig. 3A). In general, we observed more associations with glucose metabolism than insulin levels, suggesting that the microbiome in our cohort was linked more to glucose than insulin levels. Escherichia showed by far the most associations with T2D measures and notably associated with all glucose measures included in the study. Given the observed genus-specific patterns of association with T2D this raised the question how one could identify a subset of genera that were consistent markers of overall disease progression.

**Figure 3.**
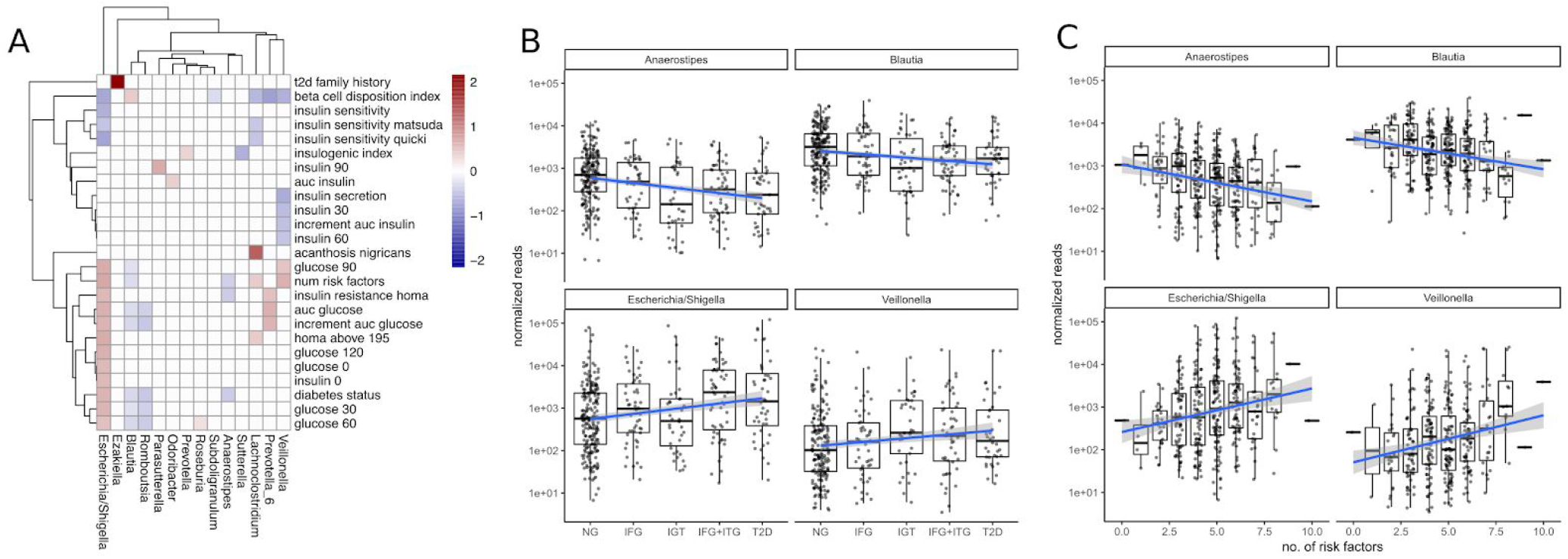
Associations between the microbiome and disease progression. (A) Significant associations (FDR < 0.1) between bacterial genera and T2D clinical variables. White boxes denote lack of significant associations and fill denotes coefficient of association between genus and variable (log2 fold change in genus abundance if the variable is increased by one standard deviation). (B) Associations between disease state and selected bacterial genera. Blue lines indicate regression lines and light gray bands denote the standard error of the regression. (C) Overall T2D risk was evaluated by the number of T2D risk factors which associated with the same genera as observed earlier. This relationship was gradual across the number of risk factors.

### A group of distinct bacteria mark the gradual progression of type 2 diabetes

To identify bacterial genera that were strong markers for disease progression we asked whether some of the 18 genera associating with diabetes measures would do so in a gradual manner across disease progression and risk. Disease progression was quantified by ordering the metabolic groups by severity ranging from normoglycemic (NG) to fully developed T2D. Disease risk was assessed by a set of manually chosen binary indicators (absent/present) for known risk factors and counting their occurrences for each individual (see Materials and Fig. S2). Thus, an individual with 8 risk factors would be considered at higher general risk for developing T2D than an individual with only 2 risk factors. Metabolic groups and the number of risk factors did only moderately correlate with each other (Spearman rho=0.45), confirming that they described different aspects of the disease. Treating the metabolic groups as well as the number of risk factors as continuous descriptors we identified a set of 4 bacterial genera that associated at least weakly with both of them (Escherichia, Veillonella, Blautia and Anaerostipes, FDR-corrected p<0.1).

We found that Escherichia and Veillonella were positively associated with diabetic state, increasing in abundance with disease progression from normal to T2D (Fig. 3B). Conversely, Blautia and Anaerostipes abundances declined with disease progression (Fig. 3B). Whereas Escherichia and Veillonella are both associated negatively with alpha diversity (Shannon index), Anaerostipes and Blautia did not (compare Fig. 2C). Therefore the protective association between these genera and T2D cannot be explained by an increased diversity alone. Intriguingly, more than 99% of the Anaerostipes sequence variants with unique species assignments belonged to the species *Anaerostipes hadrus*, a known butyrate producer. The 4 identified genera genera showed a continuously increasing or decreasing trend with disease progression, with only the prediabetes group (IGT) showing some deviation from this trend (Fig. 3B).

For all of the identified genera the number of risk factors aligned linearly with the log-transformed counts. Median Escherichia levels increased by almost 2 orders of magnitude between individuals with 2 and 8 risk factors respectively and Anaerostipes decreased by one order of magnitude (Fig. 3C). Notably, individual binary risk factors did show only very few associations with the identified genera (Fig. S4). Thus, the accumulation of T2D risk factors across the entire cohort, including healthy individuals, itself is gradually linked to changes in the microbiome.

All of the 4 presented genera also associated with the primary clinical indicators for T2D. Higher levels of Escherichia and Veillonella accompanied higher area under the glucose curve, diminished beta cell function and lower insulin levels (Fig. 4). However, Escherichia was the only genus that significantly associated with glycated hemoglobin (log2 fold change 0.5, FDR-adjusted p=0.04). Higher levels of Blautia and Anaerostipes on the other hand associated with lower area under the glucose curve, normal beta cell function and higher insulin levels (Fig. 4). Thus, the associations with markers of metabolic health were consistent with the results from oral-glucose tolerance tests.

**Figure 4.**
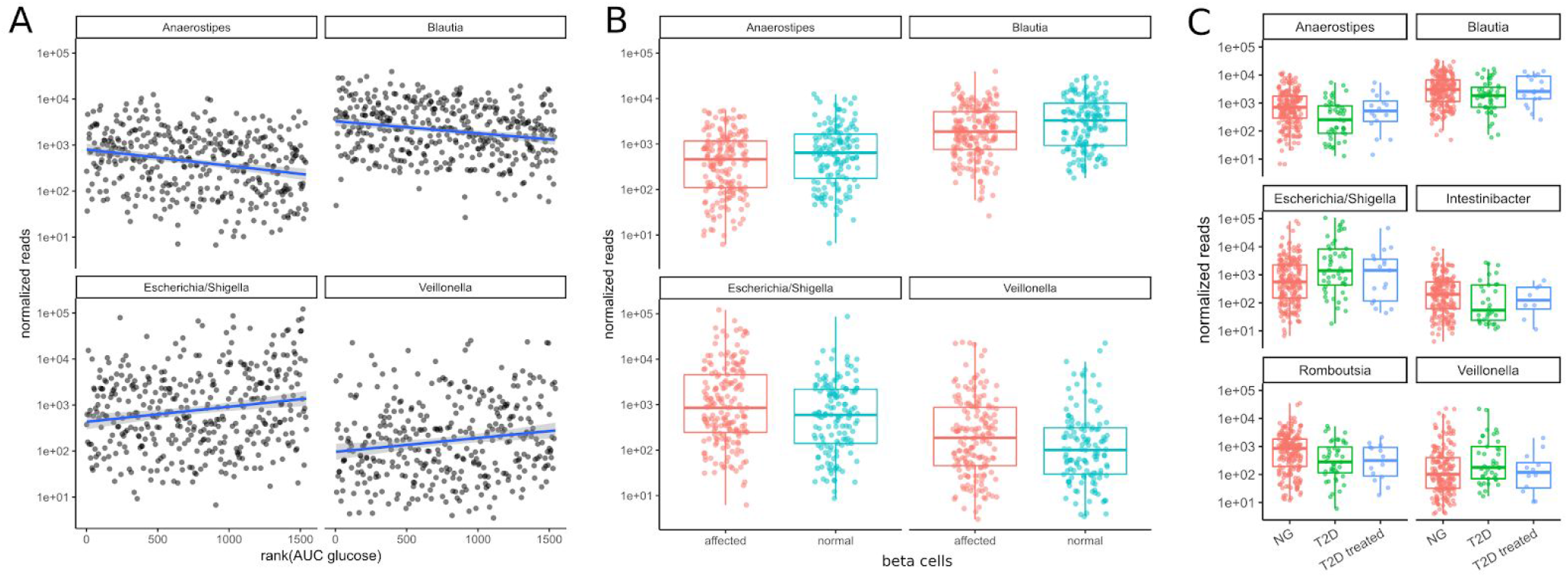
Associations between bacterial genera and the primary T2D-related clinical measurements. (A) The identified genera associated with the area under the glucose curve (AUC glucose). AUC values were rank-transformed in order to make the regression independent of outliers. The blue line denotes a linear model between log-transformed normalized counts and rank transformed AUC values. (B) Bacterial abundances stratified by beta cell function (“affected” meaning beta cell function was negatively affected). Normal beta cell function was identified by a beta cell disposition index larger than 2 (see Fig. S1B). (C) T2D treatment restored some of the altered bacterial genera (Anaerostipes, Blautia, Escherichia, Veillonella) to their normal levels but this was not true for all of them (Romboutsia remained at low levels). Mann-Whitney p-values <0.02 for all NG vs. T2D comparisons and >0.4 for all T2D vs. T2D treated comparisons except Romboutsia (p = 0.07).

We then asked whether the patterns of these 4 microbiome markers of disease might be reversed by treatment. In a control group of subjects that had already received T2D treatment, we noted that type 2 diabetes treatment (mostly metformin alone or in combination with other drugs) led to an approximate return of the 4 genera to normal levels (Mann-Whitney p values between 0.4 - 0.9, see Fig. 4C). This behavior was not observed for all genera. For instance, Romboutsia levels were not affected as strongly by diabetes treatment (p=0.07, Fig. 4C). Thus, anti-hyperglycemic treatment for glucose control was sufficient to return the identified genera close to normal levels and this is not the case for all bacterial genera.

### A confounder analysis across variable classes identifies diabetes-specific associations

As mentioned before, T2D shows comorbidity with many other clinical conditions such as obesity and cardiovascular disease. For instance, we observed correlations of the major glucose metabolism measurements such as the area under the glucose curve and insulin sensitivity with obesity related variables such as BMI, visceral fat and waist-to-hip ratio (see Figure 1D). Thus, there was a possibility that our observed changes across disease progression were driven by other covariates. For instance, the association between a bacterial genus and glucose metabolism might actually be a consequence of obesity which itself is associated with higher glucose levels. This is commonly known as confounding and obesity would be the confounder in that case.

To assess those putative confounding effects, we selected three groups of primary clinical variables which were available for the majority of the samples for T2D, obesity, and cardiovascular health, respectively (see Materials and Methods). Representative clinical variables were chosen by considering only variables measured for the majority of individuals (not all individuals provided information on all measures) and that showed the strongest association with bacterial abundances by themselves. For each of the previously identified bacterial genera and each variable in the 3 groups we then ran association tests with either only sex as the confounder (“without”) or with sex and all major variables from the other groups as confounders (“with”). The strength of confounding was evaluated by looking for changes in the regression coefficient for the association between the bacterial abundance and the respective clinical variable. If the coefficients were stable across the non-confounded (“without”) and (“with”) group we judged the association robust, whereas a coefficient closer to zero in the confounded setting (“with”) would indicate a diminished association when correcting for additional covariates and, thus, a spurious association.

Coefficients for the diabetes related clinical variables were not significantly impacted by the introduction of the additional confounders (see Fig. 5), whereas the coefficients for obesity related variables were almost completely abolished by adding the additional confounders. This means that the associations between the four identified bacterial genera and obesity-related clinical variables were essentially lost when correcting for diabetes status. Thus, diabetes measures explained most of the associations between bacterial abundances and obesity but not *vice versa.* Cardiovascular health was also confounded heavily by the T2D-related variables. In particular, we observed that association coefficients between the tested microbial genera and BMI, body fat, or diastolic pressure changed sign when correcting for secondary clinical variables (Fig. 5). This indicates that non-corrected associations can misinterpret the isolated effect of those clinical variables. Those spurious associations with obesity or cardiovascular disease could be observed with all of the 4 genera identified in our previous analysis. Here, only Veillonella showed residual associations with body fat and blood pressure after correction for some of the clinical variables (body fat and blood pressure) which led us to hypothesize that Veillonella seems to associate unspecifically with a variety of “bad health” markers.

**Figure 5.**
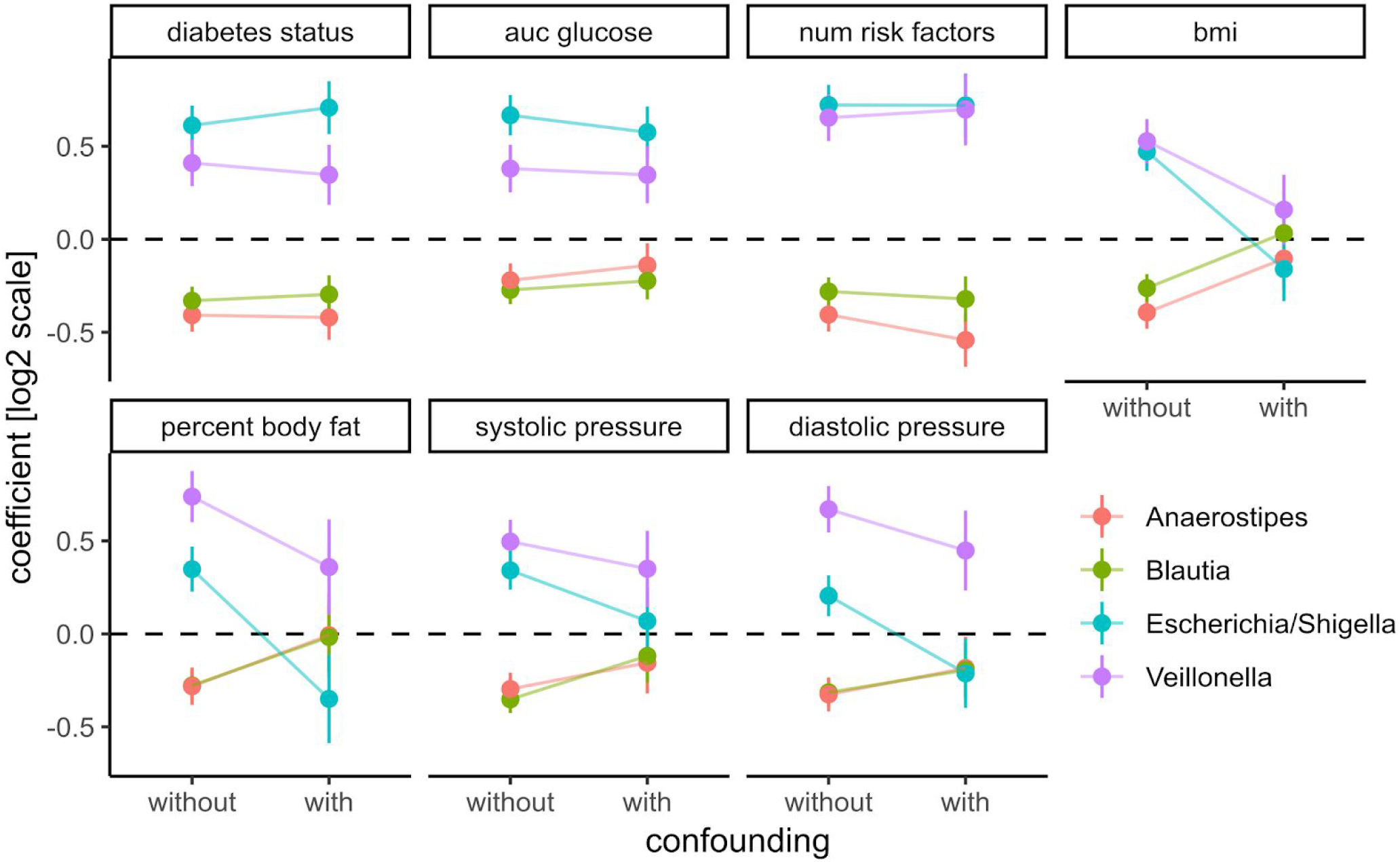
Adding prominent confounders from other classes of clinical variables did not influence effect size for diabetes related clinical response variables but did abolish associations in obesity and some cardiovascular responses. Clinical variables are grouped into T2D, obesity, and cardiovascular disease and association tests between each bacterial genus and variable are either not confounded with additional variables (without confounding) or confounded with all variables from the other groups (with confounding). Points denote the coefficient associated with the response variables under the DESeq2 model (log fold change associated with an increase of one standard deviation in the clinical variable) and error bars denote the standard errors of the model coefficient. Colors denote bacterial genera.

## Discussion

One of the challenges in studying the connections between the gut microbiome and T2D is the strong effect of medication on the gut microbiota. Metformin in particular has been shown to induce changes in the microbiome that may themselves alleviate some of the symptoms of T2D either directly or indirectly^22^. Consequently, T2D medication with metformin may mask T2D-specific changes in microbial composition. We confirm this in our study and avoided those treatment-specific effects by concentrating on a large treatment naive cohort. This allowed us to identify a set of four bacterial genera that are closely connected to T2D disease progression and risk in treatment naive individuals of a high risk population. Notably, all of the four identified genera returned to near normal levels in treated individuals. Consequently we found that metformin does not only affect more taxa in the gut microbiome than suggested previously, but may also completely disguise microbial changes induced by T2D (Fig. 4A). It is unclear whether this medication-induced restoration of the gut microbiome is a consequence of alleviated symptoms such as the regulation of blood glucose levels or a direct interaction between drugs and the microbiome. However, our observation that metformin treatment counteracts microbial changes associated with T2D but not other bacteria seems to suggest that this happens in a disease-dependent manner.

Additionally, the inclusion of a complete characterization of individual phenotypes uncovered the complex pattern of connections between microbial taxa and T2D. Most (25/31) of the diabetes-related covariates included in the study did associate with at least one microbial taxa. However, individual taxa would usually associate with a specific set of clinical measurements. For instance, even though Escherichia and Veillonella both increased with disease progression, Escherichia preferably associated with measures of blood glucose whereas Veillonella associated with more insulin-related measures (Fig. 3A). Additionally, we also found that Blautia and Anaerostipes did not only decrease with disease progression but also associated with improved beta cell function and insulin efficiency, which is to our knowledge the first time this connection has been described.

We also studied the relationship between the identified bacterial genera and T2D risk based on several established T2D risk factors. Here, we found a clear pattern of microbial shift associated with the accumulation of risk factors. This complements, previous studies have described a connection between the microbiome and the coincidence of T2D diabetes but not on T2D risk itself ^23^. We observed that this association was stable even in individuals with a low number of risk factors. This is consistent with the pathophysiology of T2D and shows that T2D-specific changes in the microbiome may precede observable symptoms ^30,31^.

Deep clinical phenotyping also allowed us to control for many of the known comorbidities of T2D and confirm the robustness of our findings. For instance, we show that the strongest associations between the microbiome and obesity-related clinical indicators (BMI and visceral fat) are completely confounded by diabetes covariates and can not be maintained when controlling for diabetes status. The implications of this observation go beyond this study and demonstrate a potential for extensive confounding in microbiome-obesity studies. As we have shown, this can be avoided by extensive phenotyping of the study subjects and can help to identify effects that are specific to the studied condition and not a secondary effect of another phenotype. In particular we feel that the combination of correcting for additional phenotypes combined with studying microbial changes that are reversed by treatment are a feasible strategy to constrain the number of associations and identify connections between disease and the microbiome that are good candidates for causal relationships.

On a coarse level our study is in agreement with previous T2D microbiome studies which mostly report a depletion of butyrate producers. On a fine level however, we find that the identified genera in our study differ from what has been found in previous studies. For instance we do not find a depletion of the butyrate-producing Roseburia, Faecalibacterium or Eubacterium ^18^ but rather observe a decrease in *Anaerostipes hadrus*, another known butyrate-producer ^32^. Some studies have also reported an increase of *E. coli* ^15,16^, however we do not observe an increase in Lactobacillus or Streptococcus. Consistent with previous findings in treatment naive subpopulations, we found that T2D could only be weakly predicted from microbiome composition when correcting for metformin treatment ^23^. Hyperglycemia itself has been shown to increase the risk for enteric infection by driving intestinal barrier permeability which is consistent with the tight association we observe between Escherichia abundance and blood glucose levels ^33^. Functionally, many of the observed associations point towards gut inflammation. Blooms of proteobacteria, like E. coli, have been associated previously with an inflamed gut and are often observed in irritable bowel disease ^34,35^. Loss of Blautia has also been associated with an inflamed gut in Crohn's disease and other clinical conditions ^36,37^. Additionally, alterations in solute carrier expression as present in the Mexican population ^12^ have been observed in the development of irritable bowel disease and have been linked to inflammation ^38,39^.

Though there is some evidence that gut inflammation may be modulated by the microbiome, it is still unclear whether one could potentially target T2D via altering the gut microbiome ^40,41^. However, the observed compositional changes consistent with inflammation might be useful as markers for long-term effects of diabetes-induced phenotypes. For instance, the gut microbiome may help to identify diabetes patients with a high risk for irritable bowel disease or colorectal cancer ^42–44^. In the end, additional studies will be required to elucidate the causal connections between the gut microbiome and T2D.

## Material And Methods

### Study Population

A cross-sectional analysis was performed in patients from Guanajuato, México, from January 2015 to December 2016, as part of the University Cohort Project CARE-In-DEEP Study (Cardiometabolic Risk Evaluation and Interdisciplinary Diabetes Education and Early Prevention). For this particular study 470 participants who had anthropometric, nutritional, biochemical and metabolic evaluation, as well as a stool sample collection, were included; at the end we had complete data and microbiome composition only for 427. Based on the oral glucose tolerance test, individuals were stratified into normal glucose metabolism (*NG*, fasting glucose less than 100 mg/dl and 2h post-OGTT glucose less than 140 mg/dl), isolated impaired fasting glucose (*iIFG*, fasting glucose 100-125 mg/dl and 2h post-OGTT glucose less than 140 mg/dl), isolated impaired glucose tolerance (*iIGT*, fasting glucose less than 100 mg/dl and 2h post-OGTT glucose between 140-199 mg/dl), impaired fasting glucose plus impaired glucose tolerance (*IFG+IGT*, fasting glucose between 100-125 mg/dl and 2h post-OGTT glucose between 140-199 mg/dl), and T2D (T2D, fasting glucose more than 125 mg/dl and/or 2h post-OGTT glucose higher than 199 mg/dl). A survey was applied to collect general information about use of medications, family history, risk factors and previous diseases. The University Research Council evaluated and approved the study protocol. All participants signed an informed consent.

### Anthropometric Measurements

Weight was measured while participants were barefoot and wearing minimal clothing. Height was obtained while the participants were standing barefoot with their shoulders in a normal position. BMI (kg/m2) was obtained from standardized measurements of weight and height and was computed as a ratio of weight (kg):height squared (m2), defining normal weight when BMI was between 18.5 – 24.9 kg/m2, overweight when BMI was between 25-29.9 kg/m2, and obesity when BMI was ≥30 kg/m2. Waist circumference was measured at the high point of the iliac crest at the end of normal expiration to the nearest 0.1 cm. Body composition was assessed with electrical bioimpedance through a Tanita Scale SC-240. All measurements were performed by personnel trained to use standardized procedures and reproducibility was evaluated, resulting in concordance coefficients between 0.88 and 0.94.

### Nutritional and Physical Activity Evaluation

A validated semi-quantitative food frequency questionnaire (FFQ) was applied to evaluate dietary intake ^45^. This questionnaire included data regarding the consumption of 116 food items. For each food, a commonly used portion size (e.g. 1 slice of bread or 1 cup of coffee) was specified on the FFQ and participants reported their frequency of consumption of each specific food over the previous year. The PA level of participants was assessed using a self-administered questionnaire that was verified when the patient assist for the metabolic evaluation. The questionnaire has a validated Spanish translation ^46^, which has been adapted for use in the Mexican population. The questionnaire is self-administered and estimates the minutes devoted to the practice of different recreational physical activities during a typical week in the last year (including walking, running, cycling, aerobics, dancing, and swimming as well as playing football, volleyball, basketball, tennis, fronton, baseball, softball, and squash, among other activities). Each item includes time intervals that allow participants to detail the exact number of minutes or hours they dedicate to each form of recreational PA, as well as the intensity of each PA (light, moderate, vigorous). The total duration of each recreational PA was expressed in minutes per day. We calculated the number of hours per week devoted to each activity, which were then multiplied by the intensity of each activity, defined as multiples of the metabolic equivalent (MET) of sitting quietly. We used the Compendium of Physical Activities to assign METs to each activity ^47^

### Metabolic evaluation and oral glucose tolerance test (OGTT)

All subjects were admitted to the Metabolic Research Laboratory of the Department of Medicine and Nutrition, Division of Health Sciences at the University of Guanajuato the day of the study between 7 and 8 AM, and a catheter was placed into an antecubital vein for all blood withdrawal. Subjects will not be allowed to eat or drink anything after 10 PM on the night before, until the study is completed. After the intravenous catheter was placed and the first blood sample was drawn, the patients ingested 75 grams of glucose. Plasma samples for glucose measurement were drawn at −15, and 0 minutes and every 30 minutes thereafter for two hours, glucose was measured by colorimetric glucose oxidase. Lipid levels were measured by dry chemistry with colorimetric method (Vitros 5600; Ortho Clinical Diagnostics)). According to the glucose levels at fasting and at 2 h during the OGTT, patients were classified as following: ***NG*** = fasting glucose <100mg/dl and 2h glucose <140mg/dl, ***IFG*** = fasting glucose between 100-125mg/dl and a 2h glucose <140mg/dl, ***IGT*** = fasting glucose <100mg/dl and 2h glucose between 140-199mg/dl, ***IFG+ITG*** = fasting glucose between 100-125mg/dl and 2h glucose between 140-199mg/dl, ***T2D*** = fasting glucose >125mg/dl and/or 2h glucose >200mg/dl, and ***treated T2D*** = previous diagnose of T2D confirmed by the medical record of the patients, consumption of hypoglycemic drugs and fasting glucose >125mg/dl and/or 2h glucose >200mg/dl. HbA1c was measured according to the international guidelines by HPLC in a subset of 182 patients.

Insulin during the OGTT was measured by a solid-phase, enzyme-labeled chemiluminescent immunometric assay (IMMULITE 1000 Siemens Healthcare Diagnostics Products Ltd). Area under the glucose and insulin curve were calculate by the trapezoidal rule.

Insulin resistance was calculated by the homeostasis model assessment (HOMA_IR) and insulin sensitivity (Matsuda Index) was derived from the insulin and glucose measurements from the OGTT as previously described ^48^. Insulin secretion was calculated dividing AUCinsulin_OGTT by the AUCglucose_OGTT, acute insulin response (AIR) was calculated dividing the insulin change from 0 to 30 minutes by the glucose change from 0 to 30 minutes during the OGTT; pancreatic beta cell function was estimated by the disposition index derived from the OGTT ^49^.

## Faecal Sample Collection

Faecal samples were collected from volunteers in a sterile container, each sample was homogenized and three aliquotes placed in sterile 1 ml screw-cap tubes which were stored at −80°C prior to DNA extraction.

## Dna Extraction

DNA extraction was performed using MoBio PowerSoil DNA Isolation kit (Mo Bio Laboratories, Inc. Carlsbad, USA) according to the manufacturer’s instructions with the following modifications. After add the C1 solution and mix, 25 μL of proteinase K solution were added and mixed by vortex. Samples were incubated at 65°C for 10 minutes, during the incubation tubes were mixed by inversion each three minutes. Tubes were secured horizontally in a vortex adapter tube holder, and vortexed at 3000 rpm for 15 minutes. Samples were incubated at 95°C for 10 minutes, during this time samples were mixed as mentioned above. Total DNA was eluted in 100 μL of sterile water. DNA concentration was quantified spectrophotometrically with a Qubit (Thermo Scientific, USA) and validated by Nanodrop (ND 2000, Thermo Scientific, USA).

### 16S rRNA gene amplification and sequencing

DNA templates were used in a two-step PCR method to sequence the V4 hypervariable region of the bacterial 16S rRNA gene. Fusion primers contained a sequence complementary for the v4 region, as well as Nextera Illumina adapter sequences to allow multiplexing of pooled libraries.

In the initial PCR we employing primers that were comprised of partial Nextera adapter and the V4 targeting forward or reverse primer sequence in agree with ^50^.

NEXT_16S_V4_U515_F

5’-TCGTCGGCAGCGTCAGATGTGTATAAGAGACAGGTGCCAGCMGCCGCGGTAA-3′ NEXT_16S_V4_E786_R

5’-GTCTCGTGGGCTCGGAGATGTGTATAAGAGACAGGGACTACHVGGGTWTCTAAT-3′

For each sample, we used approximately equal amounts of DNA template (up to 12.5 ng per reaction) and the reactions were carried out with a 3 minute denature step at 94°C, followed by 25 cycles of denature at 94°C for 45 seconds, annealing at 50°C for 60 seconds and extension at 72°C for 90 seconds, with a final extension at 72°C for 10 minutes. In all reactions were used 2x KAPA HiFi HotStart ReadyMix to generate the amplicons.

The amplicons were purified using Agencourt Ampure XP beads (Beckman Coulter) with a proportion 1.25x (v/v). The PCR products were checked using electrophoresis in 2 % (w/v) agarose gels in TAE buffer (Tris-acetate-EDTA) stained with SYBR Gold and visualized under UV light.

For each amplicon, a second PCR was carried out with a 3 minute denature step at 95°C, followed by 8 cycles of denature at 95°C for 30 seconds, annealing at 55°C for 30 seconds and extension at 72°C for 30 seconds, with a final extension at 72°C for 5 minutes with 5 ul of previous purified DNA template and using primers that attaches dual indices and Illumina sequencing adapters employing the Nextera XT kit. The PCR products were also purified equal to first PCR reactions and the DNA concentration of each PCR product was determined using a Qubit® 2.0 Broad Range Assay (Life Technologies™). An Agilent TapeStation (Agilent, Santa Clara, CA) with DNA High Sensitivity kit was used to verify the size of the PCR product only to 23 amplicons.

All samples were random distributed in similar proportions in five pools and then mixed in equal amounts (to 10 nM). The final concentration of each pool was again determined using a Qubit® 2.0.

Pools were diluted to a concentration of 9 pM for sequencing using 2×250 bp paired end sequencing chemistry v2 on an Illumina MiSeq platform. All samples were distributed according to the consecutive number assigned by the experimental laboratory in similar proportions in five pools and then mixed in equal amounts (to 10 nM). The final concentration of each pool was again determined using a Qubit® 2.0. Amplicons were denatured with 0.2 N NaOH and further diluted according to the MiSeq user guide, then combined with denatured PhiX control library. PhiX was spiked into the amplicon pool at 10% relative concentration. Image processing and base calling was performed on the BaseSpace cloud from Illumina (http://basespace.illumina.com).

### Processing of 16S sequencing data

Demultiplexed MiSeq FASTQ files were analyzed using the DADA2 workflow ^51^. High read quality was ensured by filtering and trimming the reads prior to further processing. In brief, the first 5’ 10bp of all reads were trimmed and reads were truncated on 3’ to maximum length of 240 and 200 bp for forward and reverse reads respectively as a dip in sequence quality was observed after that length.

Furthermore, all reads with more than 2 expected errors under Illumina base model were removed as well. The filtered and trimmed reads were grouped by sequencing run and error model was fit for each run separately using the DADA2 default parameters. Sequence variants were obtained for each run separately using the previously calculated error models and the dereplicated input sequences. The sequence variants and counts were then joined across all runs in a complete sequence table and de novo bimera removal was run on the entire table.

Taxonomy for the final sequence variants was called using DADAs’s RDP classifier and using the SILVA database (version 132) ^52^. Species were identified separately by exact sequence matches where possible (again using SILVA version 132). The final data set was joined with clinical metadata and saved in a phyloseq object for all downstream analysis ^53^.

In order to identify additional biases or batch effects. We checked whether particular sequence variant read counts were associated with DNA extraction order, DNA extraction date or the scientist that extracted the sample. We could not identify any bias visually and the distribution of correlations between the extraction date and individual sequence variant abundances was similar to one obtained from a random Poisson model. Finally, we also verified that there were no run batch effects by PCoA plots were we observed no particular separation of samples by sequencing run. a notebook for those quality control steps can be found in the study repository as described in “Data availability”.

### Association tests

Association tests were run using DESeq2 with some custom adjustments ^54^. First, the input count matrix was filtered by a “rule of 10” where we only tested those taxa with an average count of at least 10 reads and which appeared at least in 10% of all samples. This was necessary to avoid bimodal p value distributions in the multiple tests. The count matrix was normalized across samples using the DESeq2 size factors and the “poscounts” correction for zero read counts. All continuous clinical variables were standardized (subtraction of mean and division by standard deviation). All tests used sex as a confounding variable. Age did not show major associations with any clinical variables in this study and including it as a confounder did not have any effect. Consequently, we did not include age as a default confounder in our analysis.

Association tests were then run for all combinations between taxa and clinical variable and only for those individuals with non-missing measurements. All associations discussed in detail in this manuscript were validated manually in order to confirm the lack of extreme outliers in the scatter plots. P-values were adjusted for false discovery rate using independent hypothesis weighting in order to avoid biases for tests with low abundance taxa ^55^.

### Data availability

Raw sequencing data is provided on the sequence read archive (SRA) under the Bioproject PRJNA541332. All additional primary input files as well as intermediate files and R notebooks in order to reproduce the analysis and figures is provided at https://github.com/resendislab/mext2d. More complex functions that could be potentially useful in the analysis of other data sets are furthermore provided along with documentation in their own R package at https://github.com/resendislab/mbtools.

## Supplementary Figures

**Figure S1:**
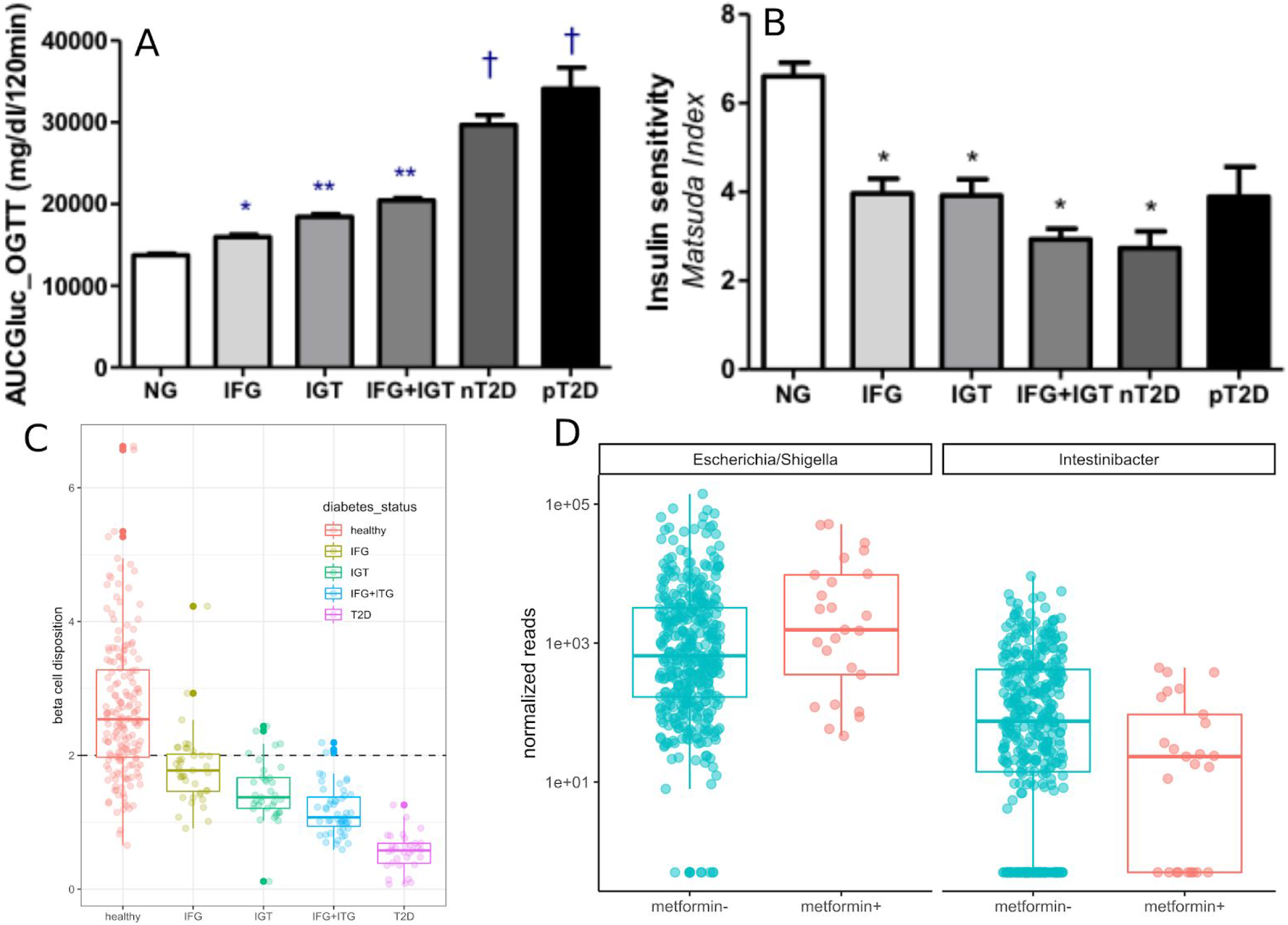
(A) Increment of the area under the glucose curve by metabolic group. “nT2D” denotes new (treatment naive) T2D and pT2D denotes previous (treated) T2D. Bars denote standard error of the mean. (B) Insulin sensitivity over the metabolic groups. Bars denote standard error of the mean. (C) Beta cell disposition index by metabolic group. The dashed line denotes the cutoff value 2 that was used to separate functional from non-functional beta cells. (D) Abundances of Escherichia and Intestinbacter stratified by metformin history. Superscripts denote the following: * - p < 0.01 vs. NG group, ** - p < 0.01 vs. NG and IFG groups, † - p < 0.01 vs all groups.

**Figure S2:**
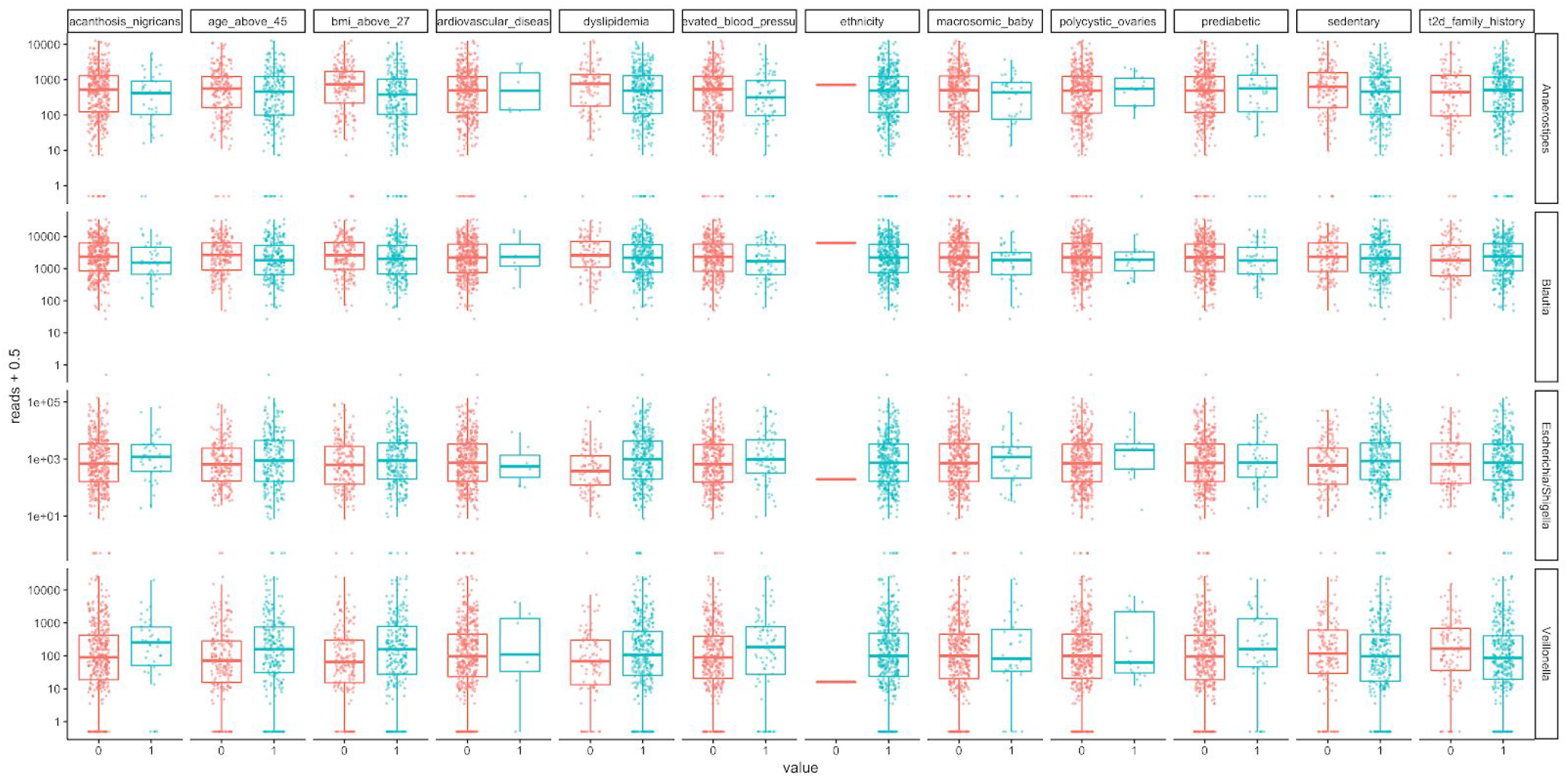
Binary risk factors used to calculate overall risk. Shown are bacterial abundances stratified by individual risk factors. Absence is denoted by zero and presence by 1. The following 3 risk factors were summarized into a single one (any of the 3 present) when calculating overall risk: polycistic ovary syndrome, cardiovascular disease, baby born with more than 4kg of weight (macrosomic). Reads were normalized across samples as described in Methods.

## Acknowledgements

The authors thank the financial support from an internal grant of the National Institute of Genomic Medicine (INMEGEN, México) and from the MIT International Science and Technology Initiatives (MISTI-2015-2016). RGM received funding from research projects 010/2014, 018/2015 and 1098/2016 from the University of Guanajuato. CMGN received a PhD scholarship from the CONACYT. We would like to thank Naeha Subramanian for valuable discussions. The authors thanks the technical support from Alfredo Mendoza and genome sequencing facility at INMEGEN, México.

## Author contributions

ORA, RGM, EA designed the study and wrote the manuscript. CD analyzed the data and wrote the manuscript. LR, LJ, MM, and CG performed extracted the samples and obtained the clinical data. NDC and VZ performed the co-occurrence and machine learning analyses and wrote the manuscript. ET analyzed the diet data. All authors discussed and proofread the final manuscript.

## Notes

https://github.com/resendislab/mext2d

https://www.ncbi.nlm.nih.gov/bioproject/PRJNA541332

